# Inhibition of cancer cells in culture. The effects of treatment with ketone bodies and/or rapamycin treatment

**DOI:** 10.1101/2021.06.27.450024

**Authors:** Anna Miller, Bo Lin, Matthew R. Pincus, Eugene J. Fine, Richard D. Feinman

## Abstract

**Background:** The potential for ketogenic diets or administration of exogenous ketone bodies to treat or prevent to cancer remains encouraging. Of particular interest is the possibility that, whatever the effect of a nutritional intervention alone, the diet might enhance the effect of existing cancer drugs, thereby requiring lower doses and a reduction in toxicity and side effects.

**Methods:** SW480, a human cell line derived from colon, was treated with ketone bodies (sodium 3-hydroxy butyrate (common name, □-hydroxy butyrate) or with sodium acetoacetate in the presence or absence of rapamycin. Cells were incubated for 96 hours in DMEM with 10 mM glucose medium. HSF2617, a human epithelial fibroblast line served as control and cells were subjected to similar treatment as the SW480 cells. Cell proliferation and glucose consumption were determined with standard reagents.

**Results:** The ketone bodies inhibited proliferation of SW480 cells in culture. Rapamycin also inhibited proliferation and its action was enhanced by the ketone bodies although there was little synergistic effect under these conditions. Human fibroblast controls were not inhibited by the ketone bodies. Both SW480 and control lines showed consumption of glucose during a 96 hour incubation period, suggesting that normal controls can switch to ketogenic metabolism while the cancer cells, which proliferate poorly, cannot. Results are consistent with recent reports of a mouse model showing the synergy of rapamycin and a ketogenic diet (Zou Y, *et al*. (2020) *PLoS ONE* **15** (5)) as well as earlier publications describing additive or synergistic effects of ketogenic diets with other modalities of cancer treatment.

**Conclusions:** The results show that the growth of a cancer cell line in culture can be inhibited by the addition of ketone bodies or rapamycin to the growth medium. The combination of treatments was found to be additive, consistent with results from a previously published mouse model. The data demonstrate the potential for a strategy whereby doses of anti-cancer agents that have detrimental or toxic side-effects can be reduced if coupled to an appropriate source of ketone bodies.

## Background

Renewed emphasis on the metabolic aspects of cancer and a likely critical role of insulin has raised the prospect of ketogenic diets as a potential cure or preventive strategy. A major theme is the role of the Warburg effect, the observation that cancer cells have a relatively greater reliance on glycolysis compared to aerobic respiration even in the presence of oxygen (Reviews: [1-5]); ketone bodies provide acetyl-CoA suggesting the possibility of by-passing whatever prevents aerobic metabolism in tumors. Beyond these effects, ketogenic diets have been shown to augment or synergize with pharmacology or other treatment modalities [6-11]. Of particular relevance to the current study is our recent demonstration that mice expressing a spontaneous breast tumor with lung metastases respond to ketogenic diet: tumor size was reduced and longevity increased. The combination of a ketogenic diet and rapamycin extended lifespan to a greater extent than either treatment alone [6]. The effects of ketone bodies and rapamycin on cultured cancer cells, reported here, are consistent with the mouse data.

## Methods

The aim of this study was to determine the effect of ketone bodies, NaAcAc and 3-HB on SW480 cells in culture and the effect of added rapamycin. Normal human HSF2617 fibroblasts served as controls.

### Cell Lines and Cultures

SW480 and HSF2617 fibroblast epithelial cells were obtained from ATCC (Manassas, Virginia). All cell lines were cultured in DMEM at 37 °C and 5% CO_2_ pressure. Medium was supplemented with 10% fetal bovine serum, 4mM glutamine, 1mM pyruvate, 10 and 25mM glucose, and 1% (v/v) penicillin and streptomycin. Cell culture reagents were from Corning through ThermoFisher.

### Chemicals and Drugs

Sodium DL-3-hydroxybutyrate and cell culture grade dimethyl sulfoxide (DMSO) were purchased from MilliporeSigma (US). Crystal violet and rapamycin were purchased from Alfa Aesar Chemicals of ThermoFisher. A stock of 1 mg/ml rapamycin was prepared by solution in sterile cell culture grade DMSO and maintenance it at -80 °C. The final DMSO concentration was maintained at 0.5% (v/v) either in the control or the treatment samples in all experiments.

Sodium acetoacetate was prepared by reaction of equmolar ethyl acetoacetate and NaOH for 12 hours at 4 °C [12]). Acetoacetate concentration was determined by a colorimetric method [13].

### Cell treatment, Cell proliferation assay and Glucose consumption assay

Cells were grown to 80% confluence and detached with trypsin-EDTA 1X solution. Cell number was determined by trypan blue stain protocol. Experimental cells were treated with chemicals and drugs at the same time in 10 mM glucose DMEM/low glucose in 96 wells tissue culture plates in parallel with controls without any additives. Crystal violet proliferation assay [14] was used to determine viable cells in all treatment conditions. Assays were measured on a Perkin-Elmer Victor3 multilabel plate reader. Cell glucose consumption was quantified by Glucose ^®^Glo assay kit (Promega).

## Results

### Ketone bodies inhibit growth in SW-480 cancer cells

SW480 cells, a line derived from colon cancers, were studied in the presence or absence of the ketone bodies. sodium acetoacetate (NaAcAc) or sodium 3-hydroxybutyrate (3-HB; common name: □-hydroxybutyrate). Proliferation after 96 hours was determined by Crystal Violet absorption at 595 nm as shown in **Figure 1**. Proliferation of cells was inhibited by the addition of NaAcAc and somewhat less effectively by 3-HB. Absorption was very low at initial seeding, typically OD_595_ = 0.2, much lower than subsequent time points, indicating that the cells were not killed by the ketone bodies but only reduced in their ability to proliferate. The primary effect of ketogenic metabolism is, in effect, the transfer of acetyl-CoA units from the liver to the periphery [1-4]. While 3-HB is the major circulating form, acetoacetate is the immediate source of acetyl-CoA. Metabolism of 3-HB involves oxidation to acetoacetate by an NAD-dependent dehydrogenase and conversion to acetoacetyl-CoA catalyzed by succinyl-CoA:acetoacetate CoA transferase (SCOT). Acetoacetyl-CoA is converted to acetyl-CoA in a thiolase-catalyzed reaction (**Figure 2)**. The apparent difference in rates of inhibition of the cells from the two ketone bodies is likely due to the dehydrogenase step becoming rate-determining. In addition, it is important to note that 3-HB mediates several other reactions, notably inhibition of histone deactylation [15]. These reactions may also have an inhibitory effect on cell proliferation but function in different cellular compartments.We have generally found that incubation with 3-HB shows greater variability and provides a somewhat reduced inhibition of proliferation as compared to acetoacetate.

**Figure 1.**
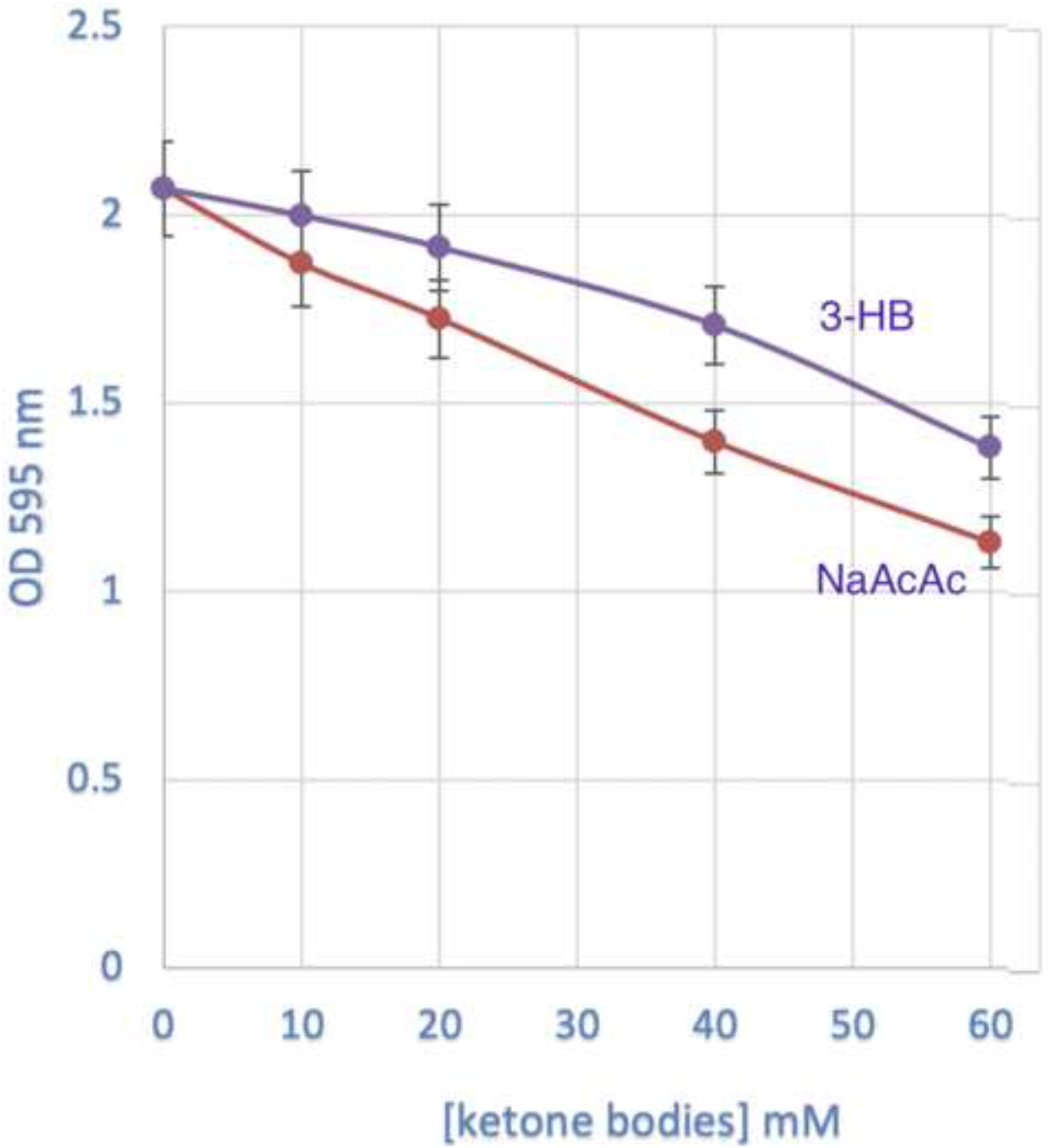
Inhibition of SW480 proliferation by ketone bodies. SW480 cells were cultured with indicated concentrations of sodium 3-hydroxybutyrate (3-HB; blue line) or sodium acetoacetate (NaAcAc; red line) in 10mM glucose DMEM 37 °C in a 5% CO_2_ incubator. Proliferation was determined at 96 hours using a crystal violet assay. (Suppl: MILLER_KB_OrigData_Figure_1.png)

**Figure 2.**
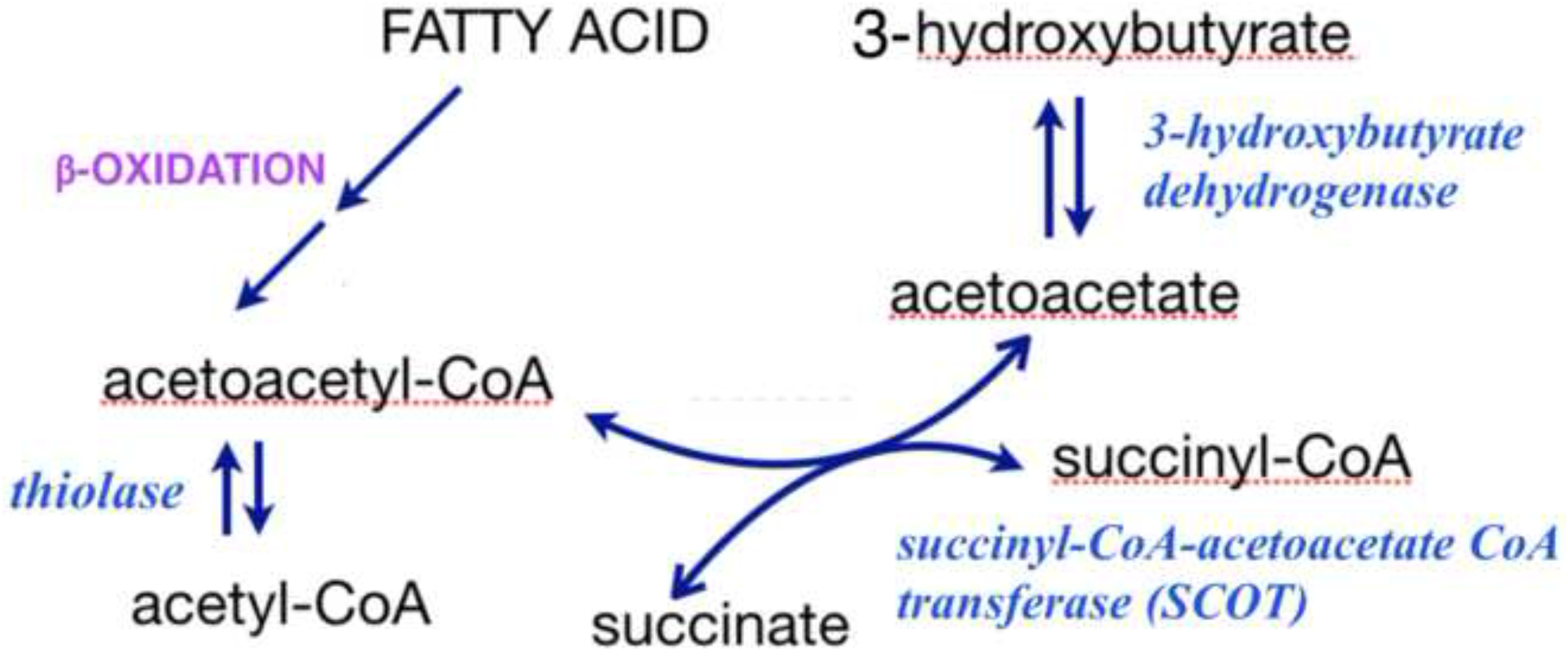
Mechanism of metabolism of ketone bodies. The dehydrogenase is an NAD-dependent enzyme and is likely rate-determining in the utilization of 3-hydroxybutyrate.

### Normal human fibroblasts are not inhibited by ketone bodies

HSF2617, a normal human fibroblast cell line, served as a control in these experiments. Cells were incubated with ketone bodies in 10 mM glucose for 96 hours (**Figure 3**). The addition of ketone bodies showed little reduction in proliferation. Micrographs of the cancer cells and controls were consistent with the specificity of inhibition (**Figure 4)**.

**Figure 3.**
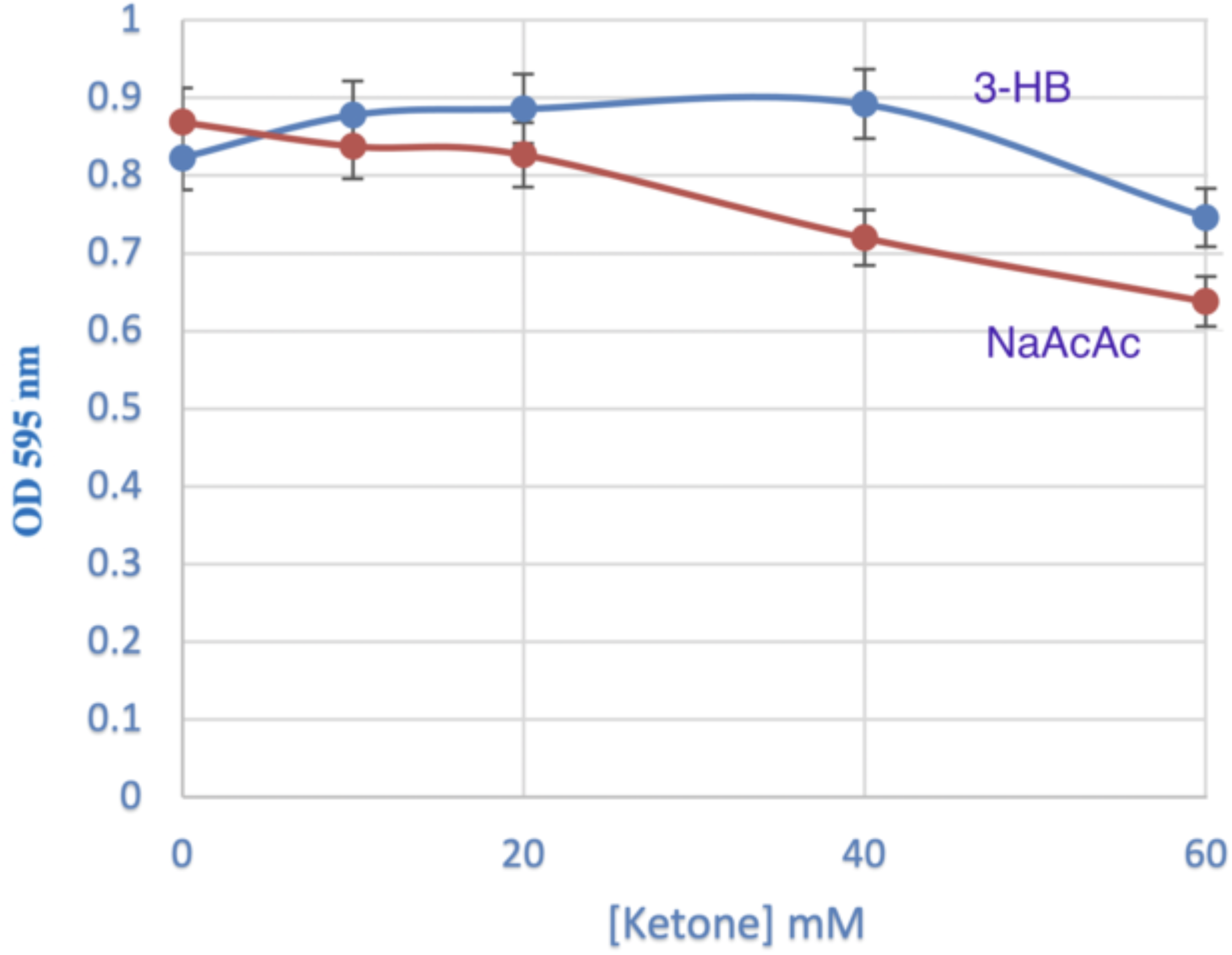
Proliferation of normal human fibroblasts is not inhibited by ketone bodies. HSF 2617 fibroblasts were cultured with indicated concentrations of 3-HB (blue line) or NaAcA (red line) in 10mM glucoseDMEM 37 °C in a 5% CO_2_ incubator. (Suppl: MILLER_KB_OrigData_Figure_3.png)

**Figure 4.**
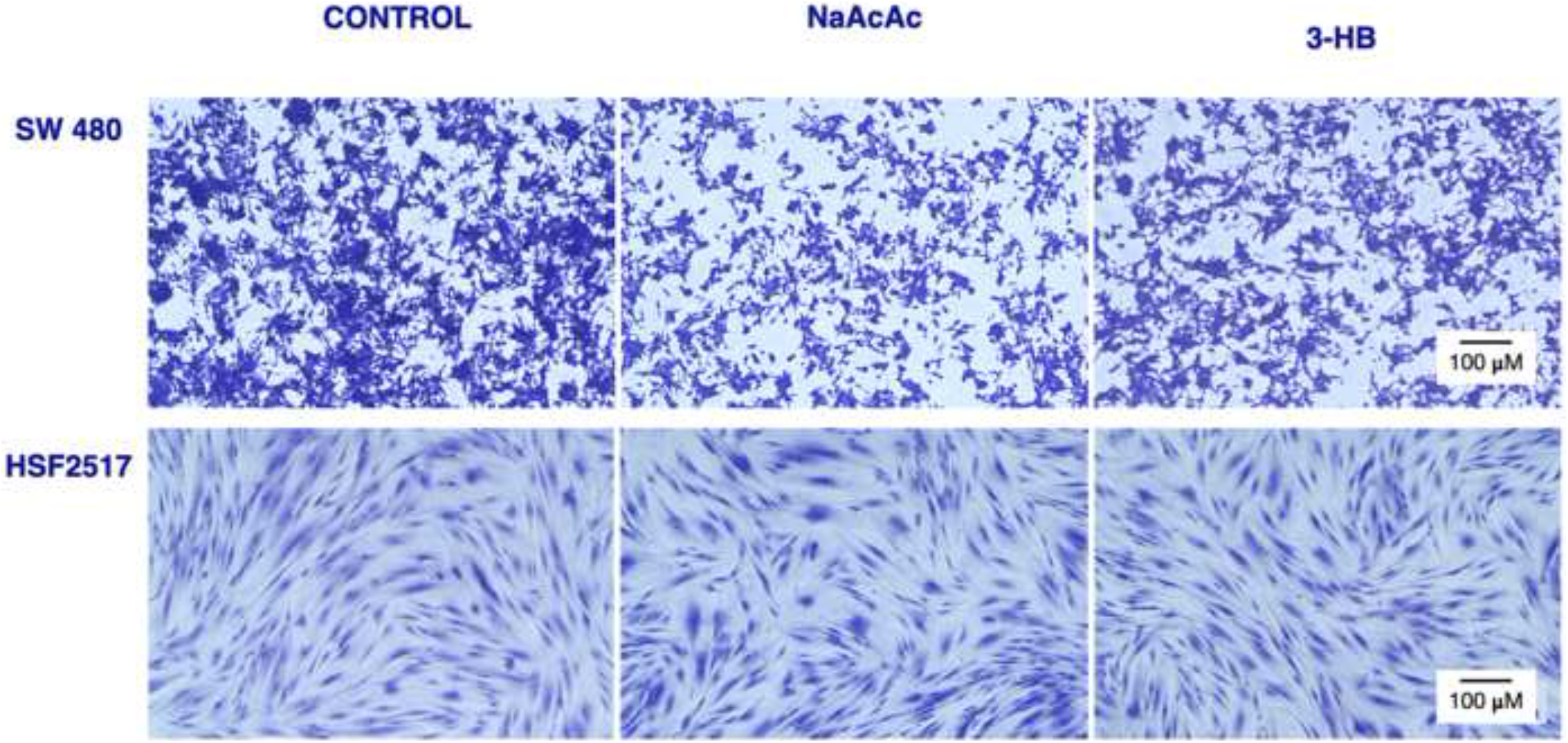
Morphology of SW480 and fibroblast controls after treatment with ketone bodies. SW480 or HSF2617 cells were cultured as described in Figures 1 and 3 and treated with 40 mM ketone bodies at 96 hours as indicated. (Original data shown with added label)

### Glucose is consumed during incubation with ketone bodies

#### Response of SW480 cells

In these studies, cells were grown in a glucose medium (10 mM glucose, DMEM) and glucose remaining in the medium over time was measured with the commercial ^®^Glo assay. The SW480 cells were incubated in the absence of ketone bodies, or in the the presence of 10, 20, 40 or 60 mM Na AcAc (**Figure 5, panel A**). In parallel experiments, the cancer cells were incubated with identical concentrations of 3-HB as shown in **panel B** of the figure. (Note that in these studies, SW480 cells show a lag in the time to adherence and the proliferation zero time is approximately 20 hours after addition of cell. This is indicated by the light blue shading in **Figure 5, panels A** and **B**). The two ketone bodies, NaAcAc and 3-HB, showed different patterns of glucose depletion of the SW480 cancer cells. NaAcAc caused a dose-dependent reduction in remaining glucose in the medium (**panel A**, red lines); the glucose half-life in the presence of NaAcAc was progressively reduced from baseline of about 55 hrs to under 40 hours at 60 nM, the highest acetoacetate concentration. In distinction, in the presence of 3-HB, cells maintained a constant concentration-independent rate of glucose utilization. None of the responses to 3-HB exceeded the baseline value found in the absence of ketone body (**panel B**, blue lines) and all four responses were superimposable. To emphasize the marked difference in effects of the two ketogenic agents, the curve for 60 mM NaAcAc was redrawn from panel A (broken gray line in **panel B)**. We interpret these data, as in the proliferation data in **Figure 1**, as a reflection of the rate-limiting effect of the dehydrogenase, that is, a slow conversion of 3-HB to acetoacetate. Involvement of 3-HB in other processes may also contribute to the inhibiting effect on proliferation and nutrient consumption.

**Figure 5.**
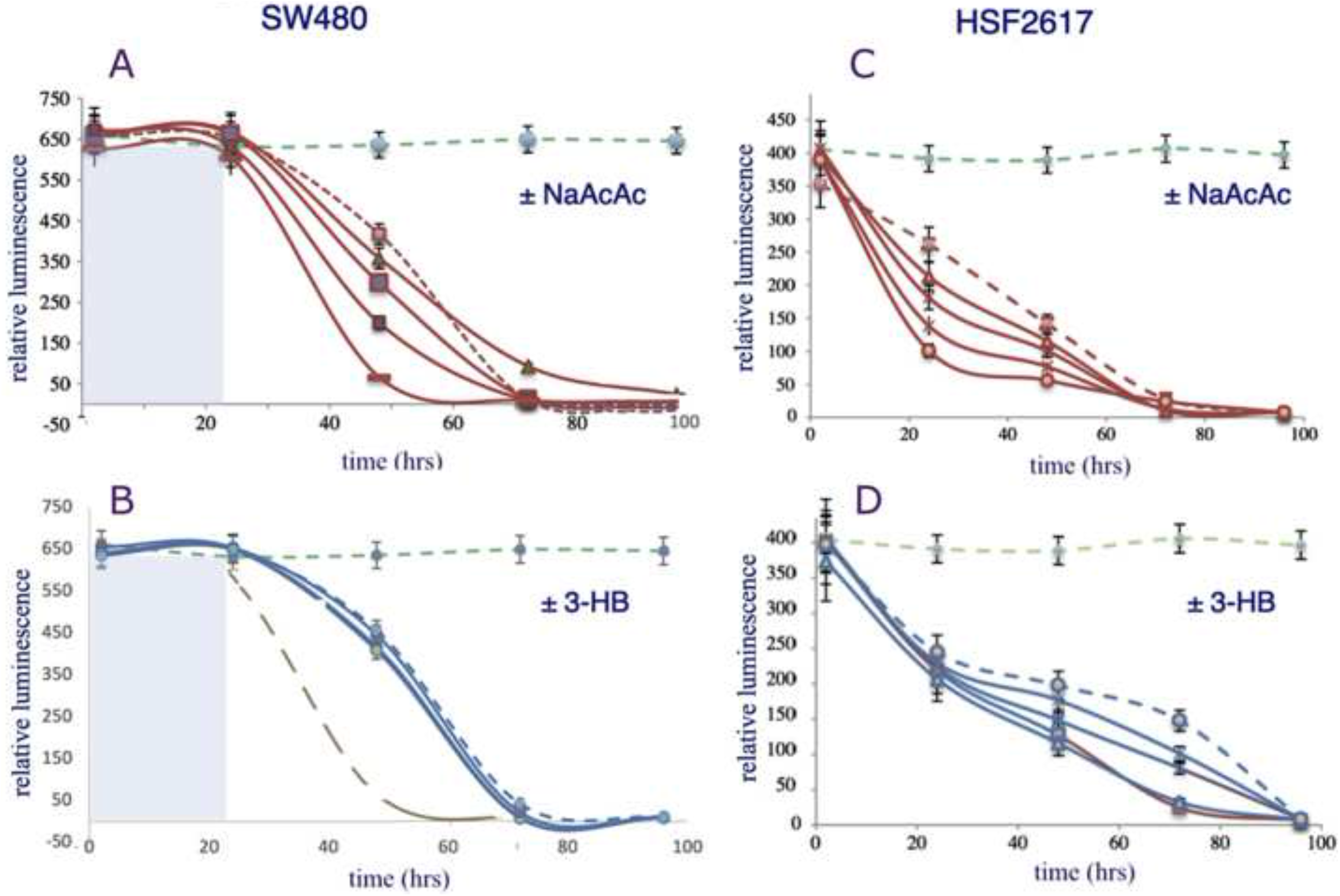
Glucose consumption and the effects of ketone bodies. SW480 cancer cells (**panels A** and **B**) or HSF2617 fibroblast controls (**panels C** and **D**) were cultured in 10 mM glucose DMEM 37 °C, 5% CO_2_ incubator. Glucose remaining in the medium was measured in triplicate at 2, 24, 48, 72 and 96 hours using the ®Glo relative luminescence assay. (Suppl: MILLER_KB_OrigData_Figure_5_A_B.png and MILLER_KB_OrigData_Figure_5_C_D.png) Cells were cultured without ketone bodies (broken red or blue lines) or with 10, 20, 40 or 60 mM sodium acetoacetate (NaAcAc; **panels A** and **C**, red lines) or sodium 3-hydroxybutyrate (3-HB; **panels B** and **D**, blue lines) as indicated by decreasing values at each time point. Green broken lines represent medium with no cells. In panel B, broken gray line represents the response to 60 mM NaAcAc that is redrawn from **panel A f**or comparison to the response to 3-HB. Light blue area: period of attachment of cells which represents, T = 0 approximately 20 hours.

### Response of HSF2617 human fibroblast control cells

Control HSF2617 human fibroblasts were treated with the same sequence of ketone bodies as the SW480 cells. **Figure 5** indicates that the effect of NaAcAc (**panel C**) was somewhat different from the 3-HB effect (**panel D**) but both responses were different in detail from the corresponding effects of the two ketone bodies on SW480. After about 60 hours, the HSF2617 fibroblasts still have significant remaining glucose, in distinction to the response to the SW480 cancer cells which show no remaining glucose in the medium at the equivalent time period (absolute time 80 hours). The results are consistent with the idea that the control fibroblasts are still proliferating and can switch to ketone bodies (even while glucose is still available). The cancer cells cannot effectively utilize the ketone bodies and persist in using glucose which, as it is exhausted leads to the reduction in proliferation.

One practical point is that the medium used in these experiments is higher in glucose than physiologic plasma values but it must be remembered that the uptake and biological effect of glucose in an *in vivo* system is controlled by insulin and other agents. There is no direct way to compare a plasma concentration with one in an *in vitro* preparation. The relation of the culture medium to the environment in animals or humans is therefore conjectural. While tissue culture is always a model system, it is the pattern of response that is key here. We found that 10 mM was the lowest concentration that gave reliable results.

### SW480 Cells are inhibited by rapamycin. Inhibition is additive but shows limited synergy with the effect of ketone bodies

**S**W480 cells were seeded at 8000 cells/well in 10 mM glucose in DMEM buffer and were treated with rapamycin in addition to ketone bodies for 96 hours. As shown in **Figure 6**, there is progressive reduction in cell number with increasing concentration of either NaAcAc or 3-HB. The effect of rapamycin appears to be immediate (within 24 hrs) and accounts for the initial drop in viable cells. It is noteworthy that the ability of 3-HB to enhance the effect of rapamycin is essentially the same as the effect of NaAcAc despite the difference in effectiveness of the two ketone bodies as sole agents in repressing proliferation (**Figure 1**) or in glucose consumption (**Figure 5**). The curves are roughly parallel indicating that the effects are primarily additive. The data in **Figure 6** were replotted in **Figure 7** to show the effect of concentration of ketone body on dose-response to rapamycin. If the results can be generalized to animal or human studies, the broken lines in the figure show that 20 mM acetoacetate could reduce the IC_50_ rapamycin dose, in this case, by approximately half.

**Figure 6.**
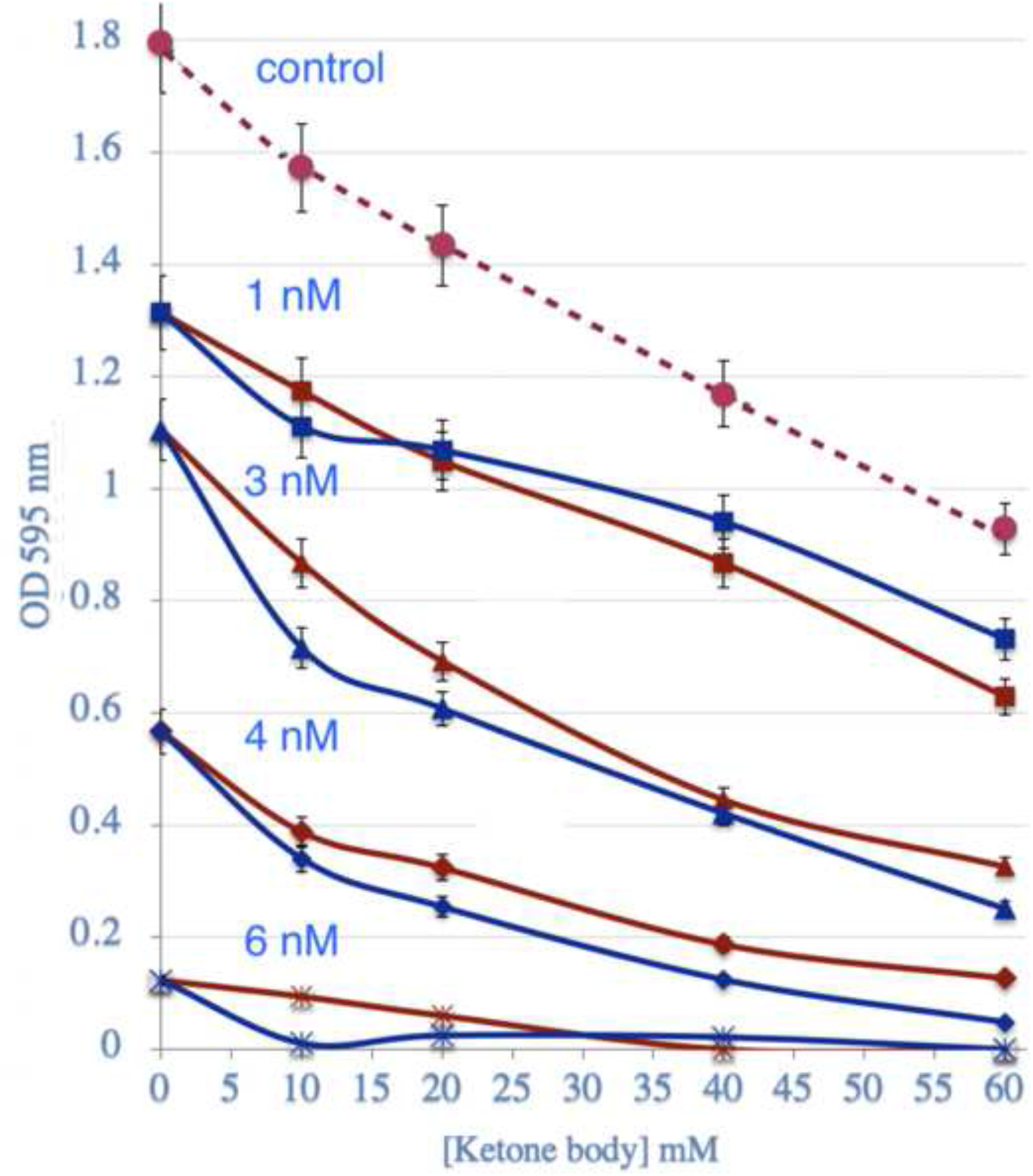
Additive effect of Rapamycin and ketone body treatments. SW480 cells were treated with sodium acetoacetate (red lines) or 3-hydroxybutyrate (blue lines). Measurements were made without rapamycin (broken green line for Na AcAc) or with addition of 1, 3, 4 or 6 nM rapamycin. Proliferation was determined with crystal violet (OD 595 nm) in triplicate at 96 hours at 37 °C in a 5% CO_2_ atmosphere. (Suppl: MILLER_KB_OrigData_Figure_6_7.png)

**Figure 7.**
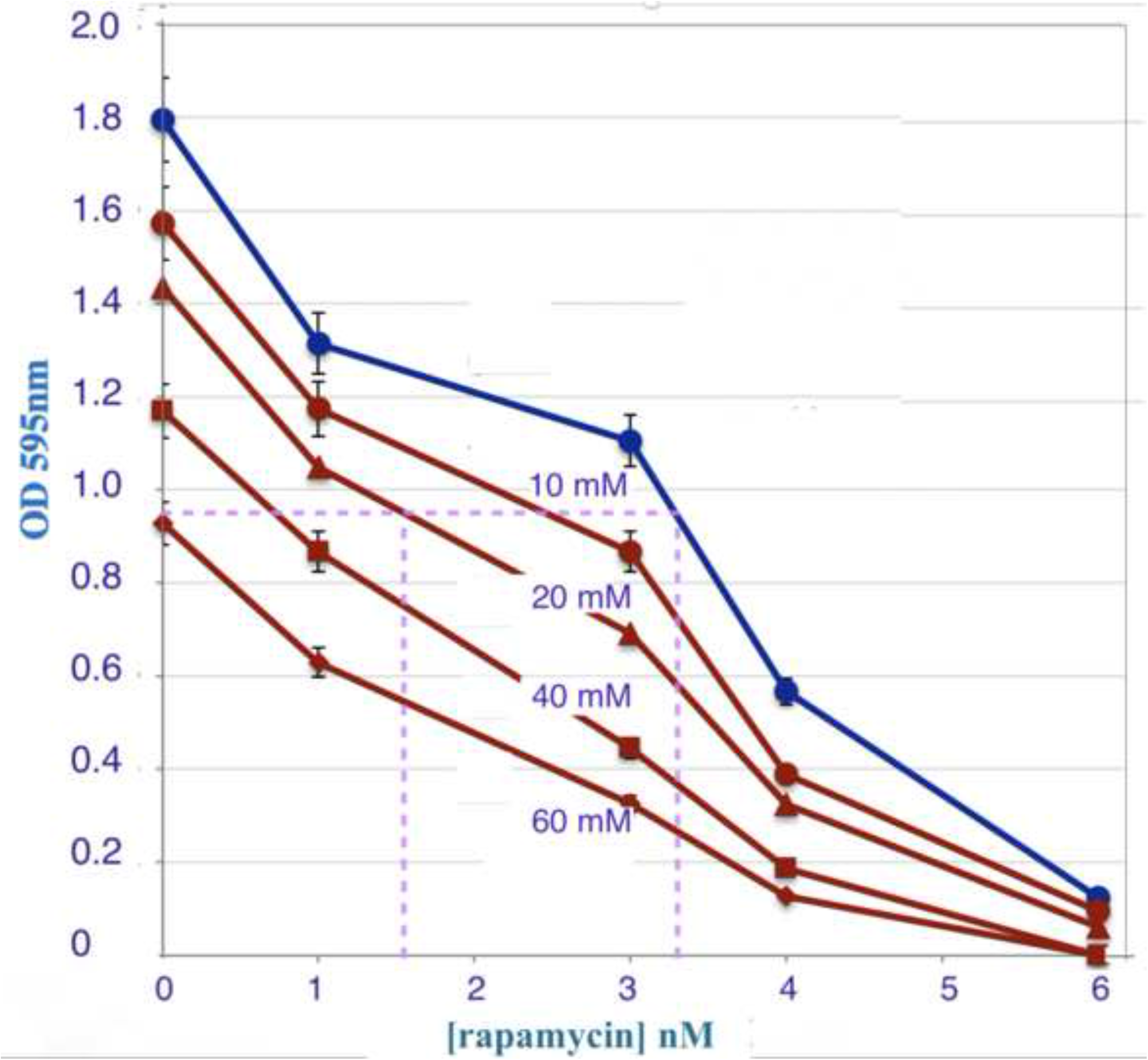
Synergy of Rapamycin and ketone body treatment. Replot of data under the same conditions as **Figure 6** (Suppl: MILLER_KB_OrigData_Figure_6_7.png)

The data in **Figure 6** were replotted in **Figure 7 for NaAcAc** to show the effect of concentration of the ketone body on dose-response to rapamycin which would be the protocol of interest from the point of view of application. For each level of rapamycin, there is a progressive decrease in OD. Inspection of the Figure shows that degree of inhibition is approximately proportional to the concentration of NaAcAc and there is no substantial synergy. This is a consequence of the rate of inhibition and the long incubation period required. There were separate effects — rapamycin rapidly inhibits cells and proliferation is inhibited more slowly by ketone bodies. To demonstrate (or exclude) inhibition, addition of the two inhibitors would have to be separated in time of administration and kinetic effects measured. Because the number of cells is very low before the addition of inhibitors and greater at early time points, the results are consistent with inhibition of proliferation rather than cell death. In the case of our mouse study, rapamycin was added against a background of chronic ketogenic diet and an analog of these conditions are currently being studies. The combined effects are consistent with the animal study. If the results can be generalized to animal or human studies, the broken lines in the figure show that 20 mM acetoacetate could reduce the IC_50_ rapamycin dose by approximately half.

## Discussion and Conclusions

Here we have shown that ketone bodies will inhibit proliferation of SW480, a colon cancer line, in culture. Control fibroblasts are not affected. From the measurements of glucose remaining in the medium, it appears that the cancer cells, as proliferation is repressed, consume glucose somewhat more rapidly than the controls, and, although speculative, we interpret this as an inability of the cancer cells to switch to ketone bodies as an alternative source as glucose is exhausted. SW480 is also inhibited by rapamycin and the combination of ketone bodies and rapamycin is additive but synergy could not be demonstrated under conditions of the experiment. The results provide an *in vitro* model for our recently published mouse study in which a ketogenic diet and rapamycin were found to be better in combination and, in this system, synergistic [6]. The conditions are not exactly the same as the animal work but the significance is that the details of the combined effects *in vitro* may serve as a guide to identify important features to explore in animals and ultimately in clinical application.

The Warburg effect casts cancer cells as lacking metabolic flexibility. In particular, entry of substrates from glycolysis or other sources into respiration or their processing in respiration is impaired. Ketogenic diets or exogenous ketone bodies may take advantage of this effect in that they by-pass the links between glycolysis and the TCA cycle, notably pyruvate dehydrogenase complex (PDH), and may relieve the limitation. Acetoacetate, directly *via* oxidation of 3-HB, provides acetyl-CoA a substrate for respiration and feedback inhibitor of PDH. In this way, the host is able to adapt to a new, or additional, fuel source. Several studies, reviewed in references [1-5] have borne out this general picture. The presumed disruption of the functioning metabolism of the cancer cell further suggests that the tumor cells may become more susceptible to other inhibitors or anti-cancer modalities of treatment. The goal is to allow lower doses of drugs or generally reduced intervention by coupling treatment with diet. Given the adverse side-effects and toxicity of most cancer drugs and the associated impaired quality of life, reducing drug dosage due to synergistic effects of diet is of evident value.

There are several examples of successful use of ketogenic diets or extrinsic ketone bodies in combination with other therapies. The results are cause for optimism but the examples are isolated and the field is in an early stage. Radiation therapy in combination with a ketogenic diet was shown to lead to permanent remission of a malignant glioma by Scheck, *et a*l. [7]. A similar synergistic effect of diet and radiochemotherapy was demonstrated by Fath and coworkers [8]. Ketogenic diets could also extend the life of mice treated with hyperbaric oxygen [9]. Drug actions that have been enhanced by diet include cisplatin [10], temozolomide [11] and 2-deoxyglucose [16]. Of note, Hopkins, *et al*. took advantage of the ability of a KD to control hyperglycemia to compensate for this effect of a PI3K inhibitor [17].

Finally, mTOR, the target of rapamycin is the subject of extensive research interest. The protein, is a kinase and senses many nutrients and has global effects, largely encouraging anabolism and proliferation (Review: [18]). Beyond the therapeutic potential of the rapamycin-ketone combination, the additive effect of ketone bodies may provide an experimental approach to dissecting the multiple inputs and functions of rapamycin. The existence of feedback loops of mTOR with the insulin pathway including a long-loop feedback inhibition of Insulin Receptor Substrate-1 (IRS-1) suggests the combination of ketone bodies and the drug may provide an experimental handle for understanding mechanism [19, 20].

## List of Abbreviations

DMSO: Dimethyl sulfoxide
3-HB: DL-3-hydroxybutyrate (commonly referred to as ⍰⍰-hydroxy butyrate)
IRS-1: Insulin Receptor Substrate-1
NaAcAc: Sodium acetoacetate
mTOR: Mammalian target of Rapamycin
PDH: pyruvate dehydrogenase complex
PI3K: Phosphoinositol triphosphate kinase
SCOT: succinyl-CoA:acetoacetate CoA transferase

## Declarations

### Ethics approval and consent to participate

Not Applicable

### Consent for publication

Not Applicable

### Availability of data and materials

The datasets used and analyzed in the current study are available from the corresponding author on reasonable request.

## Authors’ contributions

The project was initiated by EJF and RDF. AM carried out implementation of the plan, performing most of the experimental work and provided interpretation, analysis and presentation of the data. All authors participated in discussion, interpretation, revision and preparation of the manuscript.

## Acknowledgements

We are grateful to Dr. Sarah Hofmann for valuable scientific advice and information.

## References

1. Tan-Shalaby, J. Ketogenic Diets and Cancer, Emerging Evidence; Review Article. Federal Practitioner, February 2017 Oncology Special Issue.

2. Klement RJ. Beneficial effects of ketogenic diets for cancer patients: a realist review with focus on evidence and confirmation. Med Oncol. 2017;34(8):132. Epub 2017/06/28. doi: 10.1007/s12032-017-0991-5.

3. Fine EJ, Feinman RD: Insulin, Carbohydrate Restriction, Metabolic Syndrome and Cancer. Expert Rev Endocrinol Metab 2014, 9(6).

4. Poff, A., Seminars in Cancer Biology (2018), http://doi.org/10.1016/j.semcancer.2017.12.011

5. Fine EJ, Segal-Isaacson CJ, Feinman RD, Herszkopf S, Romano MC, Tomuta N, et al. Targeting insulin inhibition as a metabolic therapy in advanced cancer: a pilot safety and feasibility dietary trial in 10 patients. Nutrition. 2012;28(10):1028–35. doi: 10.1016/j.nut.2012.05.001..

6. Zou Y, Fineberg S, Pearlman A, Feinman RD, Fine EJ (2020) The effect of a ketogenic diet and synergy with rapamycin in a mouse model of breast cancer. PLoS ONE 15(12): e0233662. http://doi.org/10.137/journal.pone.0233662

7. Woolf, E.C. Syed, N. Scheck, A.C. Tumor metabolism, the ketogenic diet and beta-hydroxybutyrate: novel approaches to adjuvant brain tumor therapy, Front. Mol. Neurosci. 9 (2016) 122, http://dx.doi.org/10.3389/fnmol.2016.00122.

8. B.G. Allen, S.K. Bhatia, J.M. Buatti, K.E. Brandt, K.E. Lindholm, A.M. Button, et al., Ketogenic diets enhance oxidative stress and radio-chemo-therapy responses in lung cancer xenografts, Clin. Cancer Res. 19 (2013) 3905–3913

9. Poff AM, Ari C, Seyfried TN, D’Agostino DP. The ketogenic diet and hyperbaric oxygen therapy prolong survival in mice with systemic metastatic cancer. PLoS One. 2013 Jun 5;8(6):e65522. doi: 10.1371/journal.pone.0065522. PMID: 23755243; PMCID: PMC3673985.

10. Hsieh MH, Choe JH, Gadhvi J, Kim YJ, Arguez MA, Palmer M, et al. p63 and SOX2 Dictate Glucose Reliance and Metabolic Vulnerabilities in Squamous Cell Carcinomas. Cell Rep. 2019;28(7):1860–78 e9. doi: 10.1016/j.celrep.2019.07.027. PubMed PMID: 31412252.

11. Tieu, M. T., et al. (2015). Impact of glycemia on survival of glioblastoma patients treated with radiation and temozolomide. Journal of Neuro-Oncology 124(1): 119–126.

12. López-Soriano, F.J. and Argilés, J .M. A Simple Method for the Preparation of Acetoacetate. Fisiologia Analytical Letters, 18(B5), 589–592 (1985)

13. Schilke, R.E. and Johnson, R.E. A Colorimetric Method for Estimating Acetoacetate. American Journal of Clinical Pathology, Volume 43, Issue 6, 1 June 1965, Pages 539–543, http://doi.org/10.1093/ajcp/43.6.539

14. Feoktistova, M. Geserick, P, Leverkusold, M. Crystal Violet Assay for Determining Viability of Cultured Cells Spring Harb Protoc; doi:10.1101/pdb.prot087379 http://cshprotocols.cshlp.org/

15. Newman JC, Verdin E. Ketone bodies as signaling metabolites. Trends Endocrinol Metab. 2014;25(1):42–52. doi:10.1016/j.tem.2013.09.002

16. Marsh J, Mukherjee P, Seyfried TN. Drug/diet synergy for managing malignant astrocytoma in mice: 2-deoxy-D-glucose and the restricted ketogenic diet. Nutr Metab (Lond). 2008 Nov 25;5:33. doi: 10.1186/1743-7075-5-33. PMID: 19032781; PMCID:PMC2607273.

17. Hopkins BD, Pauli C, Du X, Wang DG, Li X, Wu D, et al. Suppression of insulin feedback enhances the efficacy of PI3K inhibitors. Nature. 2018;560(7719):499–503. Epub 2018/07/28. doi: 10.1038/s41586-018-0343-4. PubMed PMID: 30051890; PubMed Central PMCID:PMCPMC6197057

18. Hall M.N. (2016) TOR and paradigm change: Cell growth is controlled. Mol Biol Cell 27:2804–2806

19. Leontieva, O., Demidenko, Z. & Blagosklonny, M. Rapamycin reverses insulin resistance (IR) in high-glucose medium without causing IR in normoglycemic medium. Cell Death Dis 5, e1214 (2014). https://doi.org/10.1038/cddis.2014.178.

20. Yoon M. S. (2017). The Role of Mammalian Target of Rapamycin (mTOR) in Insulin Signaling. Nutrients, 9(11), 1176. https://doi.org/10.3390/nu9111176

